# Building Multivariate Molecular Imaging Brain Atlases Using the NeuroMark PET Independent Component Analysis Framework

**DOI:** 10.1101/2025.02.18.638362

**Authors:** Cyrus Eierud, Martin Norgaard, Murat Bilgel, Helen Petropoulos, Zening Fu, Armin Iraji, Granville J. Matheson, Melanie Ganz, Cyril Pernet, Vince D. Calhoun, Alzheimer’s Disease Neuroimaging Initiative

## Abstract

Molecular imaging analyses using positron emission tomography (PET) data often rely on macro-anatomical regions of interest (ROI), which may not align with chemo-architectural boundaries and obscure functional distinctions. While methods such as independent component analysis (ICA) have been useful to address this limitation, the fully data-driven nature can make it challenging to compare results across studies. Here, we introduce the NeuroMark PET approach, utilizing spatially constrained independent component analysis to define overlapping regions that may reflect the brain’s molecular architecture.

We first generate an ICA template for the PET radiotracer florbetapir (FBP), targeting amyloid-β (Aβ) accumulation in the brain, using blind ICA on large datasets to identify replicable independent components. Only components that targeted Aβ were included in this study, defined as Aβ networks (AβNs), by omitting components targeting myelin or other non-Aβ targets. Next, we use the AβNs as priors for spatially constrained ICA, resulting in a fully automated ICA pipeline called NeuroMark PET. This NeuroMark pipeline, including its AβNs, was validated against a standard neuroanatomical PET atlas, using data from the Alzheimer’s Disease Neuroimaging Initiative (ADNI). The study included 296 cognitively normal participants with FBP PET scans and 173 with florbetaben (FBB) PET scans, an analogue radiotracer also targeting Aβ accumulation.

Our results show that NeuroMark PET captures biologically meaningful, participant-specific features, such as subject specific loading values, consistent across individuals, and also shows higher sensitivity and power for detecting age-related changes compared to traditional atlas-based ROIs. Using this framework, we also highlight some of the advantages of using ICA analysis for PET data. In this study, an AβN consists of weighted voxels and forms a pattern throughout the entire brain. For example, components may have weighted values at every voxel and can overlap with one another, enabling the separation of artifacts which may coincide with the AβNs of interest. In addition, this approach allows for the differentiation, separating white matter components, which may overlap in complex ways with the AβNs, mainly residing in the neighboring gray matter.

Results also showed that the most age associated AβN (representing the cognitive control network, CC1) exhibited a stronger association with age compared with macro-anatomical regions of interest. This may suggest that each NeuroMark FBP AβN represents a spatial network following chemo-architectural uptake with greater biological relevance compared with anatomical ROIs.

In summary, the proposed NeuroMark PET approach offers a fully automated framework, providing accurate and reproducible brain AβNs. This approach enhances our ability to investigate the molecular underpinnings of brain function and pathology, offering an alternative to traditional ROI-based analyses.

## Introduction

Analysis of positron emission tomography (PET) data is usually performed based on macro-anatomical regions of interest (e.g., Desikan et al., 2006). In many cases, however, chemo-architectural boundaries do not correspond well to anatomical atlases (Beliveau et al., 2020) and even more so, they are not hard boundaries but rather spatially varying densities of molecular targets which can overlap in complex ways and vary from participant to participant. The anatomical atlas doesn’t allow voxels to be shared by multiple regions, which simplifies the anatomical representation, but may reduce the accuracy in complex areas where the boundaries of different regions might actually overlap. Here, we propose to use an independent component analysis (ICA) framework which allows for separation of the PET signal into whole brain covarying networks. ICA allows for multiple overlapping networks, across participants for a given PET radiotracer. Simultaneously, each network is kept as distinct as possible from the others, ensuring that every network remains unique. Even with unique networks, a single voxel can be part of more than one network, with each network contributing a certain “weighted intensity” to that voxel. The voxel weight represents the strength or degree to which it is associated with each network. This allows for a more nuanced understanding of how different brain networks might interact or coexist within the same anatomical space. ICA may also identify networks that are consistent across different participants, meaning the same patterns or networks are observed in similar ways across a population, which can be particularly useful in understanding common functional or molecular patterns in the brain. In PET, the molecular patterns are also affected by the radioligand used with differences in affinity and binding characteristics.

In this study we use florbetaben (FBB; Cho et al., 2020) and florbetapir (FBP) radioligands which both bind to amyloid-beta (Aβ) plaques. Aβ plaques have been shown to accumulate with healthy aging and can be measured using PET with FBP and FBB, among other radioligands (Fleisher et al., 2013; Bilgel et al., 2016; Jack et al., 2015). The following gray matter regions track both disease and healthy aging using standardized uptake value ratios (SUVR) and FBP (Fleisher et al., 2011; Fleisher et al., 2013): medial orbital frontal, temporal, anterior and posterior cingulate, parietal, and precuneus. Fleisher et al. (2013) shows that older healthy control subjects have a steeper SUVR increase after age 58 than before, while AD patients have a stronger SUVR increase disregarding age. Since FBB is similar to FBP, FBB may quantify age and disease analogous with FBP. Both FBP and FBB exhibit off-target binding in the white matter (WM) thought to be explained by their affinity for the beta-sheet structure of the myelin basic protein (Catafau & Bullich, 2015; Moscoso et al., 2022). This binding contaminates the signal originating from Aβ binding owing to partial volume effects, and affects quantification. Unlike anatomical atlases, which rely on predefined brain regions, ICA leverage the statistical independence, which can facilitate separation of different binding sources such as Aβ plaques and affinity for the beta-sheet structure into different components. This feature may be derived from ICA’s ability to separate a multivariate signal into independent non-Gaussian components or networks related to Aβ or myelin basic protein into different patterns. Therefore, ICA may account for the complex and overlapping patterns of radioligand binding, which anatomical atlases are unable to address.

A previous study closely related to ours is the blind ICA study conducted by Pereira et al. (2020), which employed the flortaucipir radioligand to identify ten tau-related independent components. Most of these components were bilateral but were not assessed for their replicability. Typically, ICA approaches are ‘blind’ and thus, in the case of spatial ICA, make no assumptions about the shape of the spatial maps. To facilitate statistical analyses, group ICA approaches were developed to provide a common inferential framework (Calhoun et al., 2001), solving component matching within a study. While this framework solves component matching within a study, comparisons across studies remain difficult, due to variations in model orders and other methodological differences that affect the extracted components. Therefore, spatially constrained ICA approaches allow a set of reference components to be used as spatial ‘priors’ which are then adapted to a given dataset (Lin et al., 2010). A leading spatially constrained ICA algorithm is the multivariate-objective optimization ICA with reference (MOO-ICAR), which performs well in capturing subject-specific information and removing artifacts (Du et al., 2016; Du & Fan, 2013). The automation of the NeuroMark pipeline was accomplished through the development of its specialized templates, notably the NeuroMark version 1 template (Du et al., 2020), NeuroMark version 2 multiscale templates (Iraji et al., 2023), and NeuroMark version 3 age-specified templates (Fu et al., 2024). The NeuroMark method circumvents manual selection of model orders and provides a framework that robustly yields reliable components across different participants and studies. This approach allows for the spatial correspondence of atlas-based approaches, which also benefits from the adaptive nature of data-driven approaches. NeuroMark has been widely applied to fMRI data and extended to structural MRI and diffusion MRI, and includes a set of modality-specific ICA templates (http://trendscenter.org/data) as well as an automated pipeline implemented in the GIFT software (Calhoun 2001; http://trenscenter.org/software/gift; Iraji et al., 2021).

For PET imaging, it is important to understand the linear relationship between SUVR (Standardized Uptake Value Ratio) and the ICA (Independent Component Analysis) loading values, as well as how SUVR affects ICA and its independent spatial components (or networks). Notably, we demonstrate that spatial ICA components are invariant to SUVR scaling. This is shown starting with the linear mixing model before SUVR, as expressed in Equation (1):

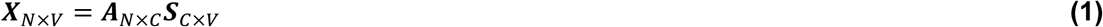

where ***X*** is the original PET data matrix for N subjects and V voxels. ***A*** is the mixing matrix, where each element in ***A*** is a loading coefficient, C is the model order (number of components). ***S*** is the source signal, containing C spatial components.

The SUVR can be conceptualized as a scaled version of the original PET signal (***X***) for each subject. Denoting the SUVR scaled signal as 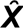, we can write 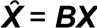, where ***B*** is a diagonal scaling matrix with subject-specific weights on the diagonal, represented as:

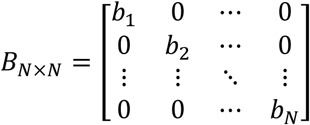

Substituting ***X*** with 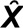, in equation 1 yields equation 2:

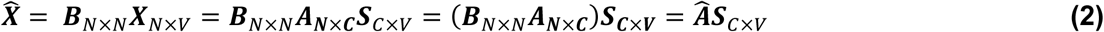

Factoring out the mixing matrix and the scaling parameter in equation 2, results in equation 3:

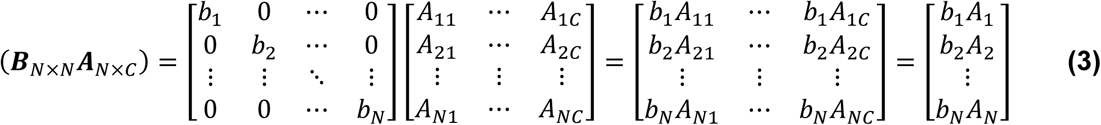

Finally, equation 2 can be simplified further into equation 4:

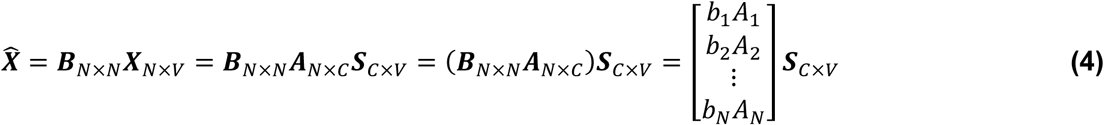

From Equations (1) through (4), we see that the source maps (***S***) are invariant to scaling, meaning SUVR scaling only affects the mixing matrix (***A***). Consequently, SUVR scaling can be applied to the loading parameters post-ICA without recomputing the components. This allows for evaluating the impact of calibration (scaling) on the results without needing to re-estimate the source components. In this study, we normalize the mixing matrix (***A***) according to each subject’s SUVR values to ensure consistency in the analysis.

In the following, we first show that blind ICA reliably decomposes PET data, here florbetapir (FBP), into biologically meaningful networks related with Aβ or myelin basic protein. Next, we generate NeuroMark ICA templates for FBP following the NeuroMark fMRI template (Du et al., 2020; Levey et al., 2022; Fu et al., 2021a; Fu et al., 2021b). Subsequently, we apply the NeuroMark template to analyze a separate dataset with florbetaben (FBB) data. Since FBB and FBP are structurally similar molecules with highly correlated binding patterns in the brain, the patterns across FBB and FBP subjects may be similar, providing a tracer-independent validation of the multivariate template. We also show through a correlation analysis between tracer binding/amyloid and age that using independent components (ICs) provide more statistical power than macro-anatomical ROIs.

## Methods

### Participants

Data used in the preparation of this article were obtained from the Alzheimer’s Disease Neuroimaging Initiative (ADNI) database (http://adni.loni.usc.edu/). The ADNI was launched in 2003 as a public-private partnership, led by Principal Investigator Michael W. Weiner, MD. The primary goal of ADNI has been to test whether magnetic resonance imaging (MRI), positron emission tomography (PET), other biological markers, and clinical and neuropsychological assessment can be combined to measure the progression of mild cognitive impairment (MCI) and early Alzheimer’s disease (AD). Details about the ADNI search criteria, resulting in 322 FBP and 198 FBB with 520 matching T1 MRI is found at the top in the supplementary methods.

We first resampled all PET images to 2×2×2 mm^3^. Then all T1-weighted MRIs (for both FBP and FBB) were processed using FreeSurfer’s recon-all function (Dale et al., 1999), resulting in a skull-stripped brain in both native and fsaverage space and cortical labeling using the Desikan-Killiany-Tourville (DKT) atlas (Desikan et al., 2006). PET frames were realigned using PETPrep_HMC (https://github.com/mnoergaard/petprep_hmc) and then registered to the T1 and MNI305 spaces using PETpipelineMATLAB (https://github.com/mnoergaard/petpipelineMATLAB). The four 5-min frames were averaged into a single PET image per participant in MNI305 space (Evans et al., 1993), and PET voxel intensities normalized to SUVR using the cerebellar cortex as the reference region as defined by FreeSurfer’s left and right cerebellum cortex. At this stage, all the DKT-ROI measures were computed by averaging voxel values in native PET space. After this, processing quality control was applied for the FBP participants, finding that the following participants needed to be excluded: 19 that failed FreeSurfer’s recon-all quality control, 2 that failed PET registration, 1 that had abnormal intensity. This leaves 300 FBP participants

Subsequently, outliers were identified and excluded based on a spatial correlation analysis. Specifically, for each participant, their spatial data was compared to the group mean. If a participant’s data differed from the group mean by more than 3 standard deviations, that participant was considered an outlier and excluded from the analysis, leaving 296 FBP participants for the remainder of the study.

For the remaining 198 FBB participants, processing quality control was applied, finding that following participants needed to be excluded: 10 that had abnormal intensities, 7 that failed FreeSurfer recon-all quality control, 5 that failed PET registration, 2 that did not fulfill spatial correlation of 3 standard deviations to mean, 1 that had a DICOM error, leaving 173 FBB participants.

Further on the 296 FBP and 173 FBB images in MNI305 space are denoted as foundational images for subsequent processing. Initially, an average SUVR map for the FBP group was created by combining all FBP foundational images. Similarly, an average SUVR map for the FBB group was generated using all FBB foundational images.

Furthermore, the foundational images derived from FBP will be categorized into groups A and B to construct the NeuroMark FBP template. Given that both FBP and FBB share numerous characteristics, including the common targeting of Aβ, the NeuroMark template, though originally developed using FBP, will be applied to FBB.

### NeuroMark Pipeline: Creating the FBP template via independent component analysis

An overview of our approach is shown in Figure 1. For the ICA processing, both FBP and FBB foundational images were smoothed using a 10 mm Gaussian kernel. Then, FBP data were split into two independent groups A and B, according to the first two rows in Table 1.

**Figure 1.**
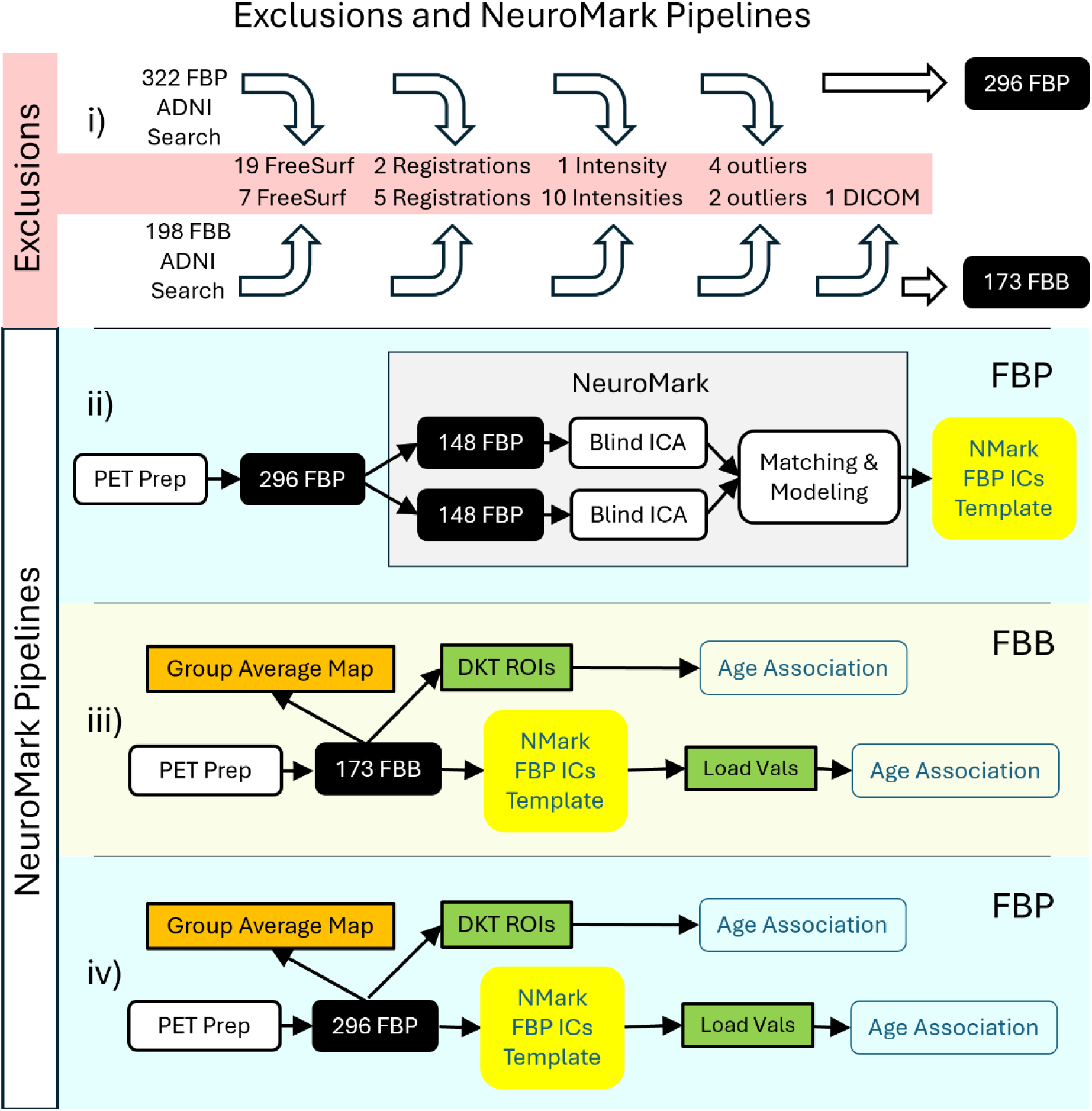
i) Depicts the exclusions (highlighted in red) before, during and after PET Prep processing until we get the foundational images in the black background boxes (more details under Participants section). ii) Depicts how the NeuroMark (NMark) independent components (ICs) FBP template is created. iii) Depicts the automated NeuroMark part generating and loading values (Load Vals) from the independent FBB data that finally is associated with age. Similarly, the average SUVR from the Desikan-Killiany-Tourville (DKT) ROIs (green background boxes) can be associated with age. In addition, mean group maps, being an average across all participants within the radioligand, are created (orange background boxes). iv) Is analogous to panel (iii) only that the FBP data was evaluated.

**Table 1.**
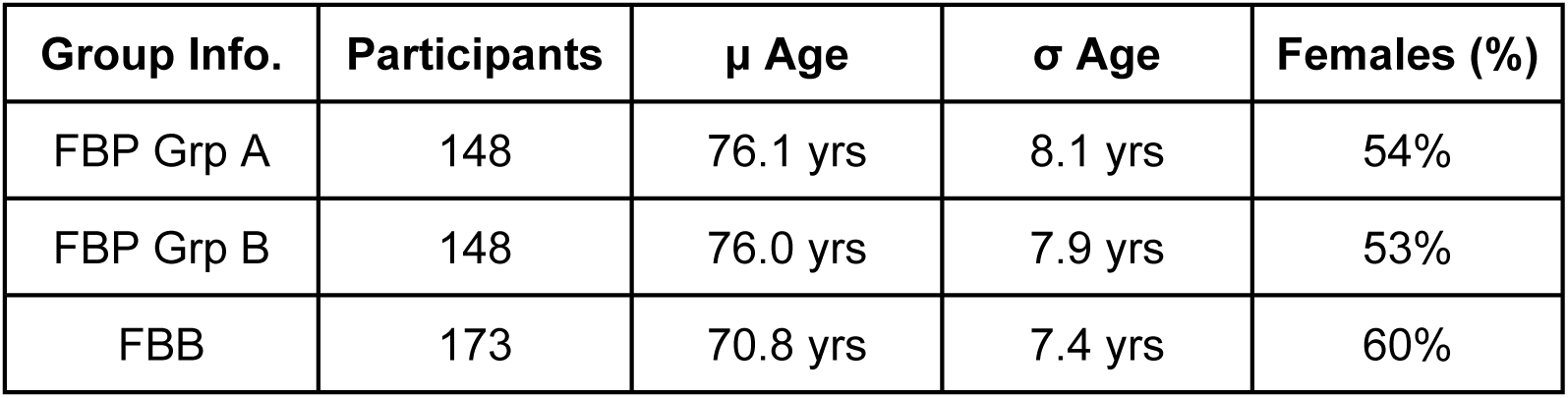
Demographics of FBP and FBB Groups (Grp), including mean (μ) and standard deviation (σ).

Spatial group ICA, using the Infomax ICA algorithm (Bell & Sejnowski, 1995), was run independently on groups A and B, using GIFT (http://trendscenter.org/software/gift; Calhoun 2001). Important ICA parameters were, a model order of 40 and no scaling of components. The 40 ICs from group A were spatially correlated with ICs from group B, and IC-pairs with a correlation below 0.4 were excluded in accordance with previous literature (Du et al., 2020). The remaining ICs represent different types of networks that were manually inspected. ICs matching white matter may be networks showing the unwanted unspecific binding to myelin basic protein, which we exclude. ICA derived networks may also represent artificial components (typically showing high values at the edges of the brain), which were further removed. Networks representing the ventricles may not be useful and were removed as well. Finally, the correlated IC pairs that remained were averaged, each creating an Aβ network (AβN) that is one part of the NeuroMark FBP template, enabling amyloid-informed constrained ICA.

In addition, we computed the participant specific loading values (explained in equation 1), similarly scaled for all n_A+B_ participants using the NeuroMark FBP template and constrained ICA on the FBP participants (n_A+B_). These loading values correspond to the degree to which a given component is expressed in a given participant.

### Evaluating the relationship between amyloid and age: Anatomical ROIs versus multivariate amyloid beta networks

After preprocessing, FBB and FBP SUVR values were calculated for anatomical ROIs in native space, using the FreeSurfer DKT atlas. In comparison, for ICA, we used MOO-ICAR together with the FBP template to calculate the loading values for each NeuroMark AβN. Next, regression analyses between age and DKT ROI SUVR, on the one hand, and between age and AβN loading values, on the other, were computed, using the simple linear regression. The regression results were then compared between DKT ROIs and NeuroMark AβNs. Significant associations are reported after false discovery rate correction (q<0.05).

## Results

### Multivariate Independent Components & Maps

The ICA processing, starting with 40 paired independent components across groups A and B, resulted in a NeuroMark template with 17 AβN. Excluded components were: 13 paired components that had lower correlation than 0.4, 4 networks that were ventricular, 2 networks that were WM related, 2 networks that represented edge artifacts, and 2 cerebellar networks that were related with the reference region, resulting in 23 exclusions. In Figure 2, the first row depicts networks from group A and the second row depicts networks from group B. The 17 NeuroMark AβNs are depicted in Figure 2 at the bottom row. Excluded networks with spatial correlation above 0.4 are displayed in the 3^rd^ column of Figure 2.

**Figure 2:**
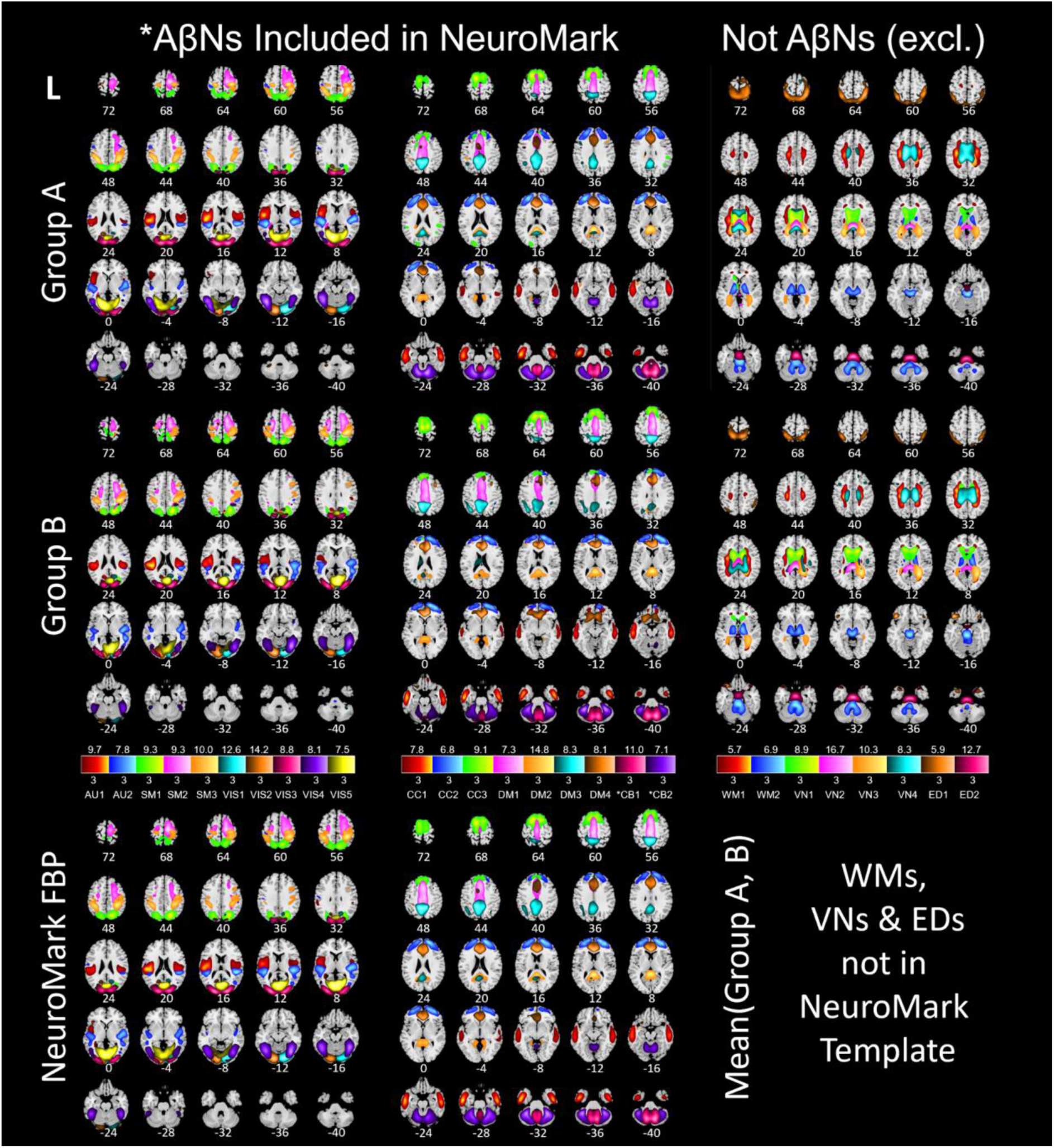
Spatial ICs from 296 FBP participants in the bottom row averaged from group A (top row) and group B (middle row). Group A (n=148) is independent of group B (n=148). All subjects are cognitively normal subjects. The color shades in the three legends correspond to the z-scores below and above each box. The shade in each color box matches voxels of its component. Acronyms: florbetapir (FBP), auditory (AU), somatosensory (SM), visual (VIS), cognitive control (CC), default mode (DM), cerebellar (CB), white matter (WM), ventricular (VN), edge (ED), left (L). The cerebellar (*CB) components are excluded due to nonspecific binding and were used as reference.

To identify the anatomical DKT region corresponding to each NeuroMark FBP IC, we matched each component to the DKT region containing the highest intensity voxel values for that IC (Table 2).

**Table 2.** Key data about AβN (excluding white matter, ventricles, edge artifacts and cerebellum). Spatial correlation between group A and group B (Corr), NeuroMark FBP Label (visualized in Figure 2), DKT ROI with NeuroMark (NMark) max point (voxel), the X,Y,Z position of the NeuroMark max intensity voxel, standardized β for FBB (explained under section “Regression of age with tracer uptake” below), FDR corrected age association (q) for FBB, standardized β for FBP, FDR corrected age association (q) for FBP. Acronyms: Negative association with age and excluded (x-neg)

| Corr | Label | DKT ROI with NMark max point | X mm | Y mm | Z mm | $\beta$ FBB | q FBB | $\beta$ FBP | q FBP |
| --- | --- | --- | --- | --- | --- | --- | --- | --- | --- |
| 0.50 | AU1 | Supramarginal Gyri | $\pm 52$ | -23 | 19 | 0.17 | 3.9E-2 | -0.078 | x-neg |
| 0.45 | AU2 | Superior Temporal Gyri | $\pm 50$ | -33 | 14 | 0.302 | 2.7E-4 | 0.051 | 0.46 |
| 0.67 | SM1 | Superior Parietal Lobules | $\pm 22$ | -73 | 45 | 0.118 | 0.15 | -0.002 | x-neg |
| 0.54 | SM2 | Precentral Gyri | $\pm 14$ | -15 | 68 | 0.169 | 3.9E-2 | 0.143 | 3.1E-2 |
| 0.52 | SM3 | Postcentral Gyri | $\pm 36$ | -39 | 53 | -0.061 | x-neg | -0.164 | x-neg |
| 0.71 | VIS1 | Right Lingual Gyrus | 16 | -89 | -15 | 0.203 | 1.8E-2 | 0.151 | 3.1E-2 |
| 0.73 | VIS2 | Left Lingual Gyrus | -12 | -91 | -15 | 0.057 | 0.53 | 0.204 | 2.1E-3 |
| 0.68 | VIS3 | Lateral Occipital Cortices | $\pm 22$ | -95 | 13 | 0.026 | 0.73 | -0.049 | x-neg |
| 0.57 | VIS4 | Fusiform Gyri | $\pm 40$ | -67 | -15 | 0.32 | 1.3E-4 | 0.147 | 3.1E-2 |
| 0.53 | VIS5 | Lingual Gyri | $\pm 6$ | -75 | 5 | 0.177 | 3.7E-2 | 0.040 | 0.55 |
| 0.52 | CC1 | Middle Temporal Gyri | $\pm 54$ | -11 | -31 | 0.428 | 6.6E-8 | 0.234 | 5.1E-4 |
| 0.62 | CC2 | Superior Frontal Gyri | $\pm 21$ | 58 | 12 | 0.151 | 6.4E-2 | 0.062 | 0.39 |
| 0.51 | CC3 | Superior Frontal Gyri | $\pm 11$ | 11 | 66 | -0.061 | x-neg | 0.118 | 7.8E-2 |
| 0.66 | DM1 | Paracentral Lobules | $\pm 2$ | 14 | 40 | 0.039 | 0.66 | 0.002 | 0.97 |
| 0.53 | DM2 | Posterior Cingulate Cortices | 0 | -43 | 9 | 0.268 | 1.4E-3 | -0.010 | x-neg |
| 0.54 | DM3 | Precuneus Cortices | $\pm 10$ | -46 | 50 | 0.234 | 5.9E-3 | 0.072 | 0.34 |
| 0.54 | DM4 | Rostral Anterior Cingulate Cortices | $\pm 5$ | 35 | 7 | 0.183 | 3.4E-2 | -0.011 | x-neg |

#### Univariate Molecular Imaging Brain Atlases of Florbetaben & Florbetapir

Next, we show an example of a Molecular Image Brain Atlas (MIBA) average from the 296 FBP subjects (Figure 3B). As expected from the literature, the WM has higher binding intensities than gray matter, at least for FBP (Nemmi et al., 2014). This non-specific binding, corresponded to two very reliable components for cortical and cerebellar WM (r=0.81 for WM1 and 0.71 for WM2). Using multiple linear regression, we estimated the weights of all 40 ICs to model the group average SUVR FBP PET. By applying these weights, we selectively subtracted the two WM and the four ventricular ICs from the group average SUVR FBP PET, revealing that the remaining SUVR is in the gray matter (Figure 3C). Figure 3A depicts that the average FBB MIBA is very similar to the average FBP MIBA (Figure 3B), bolstering that FBB may be used as test data even if the AβNs were created using FBP data.

**Figure 3:**
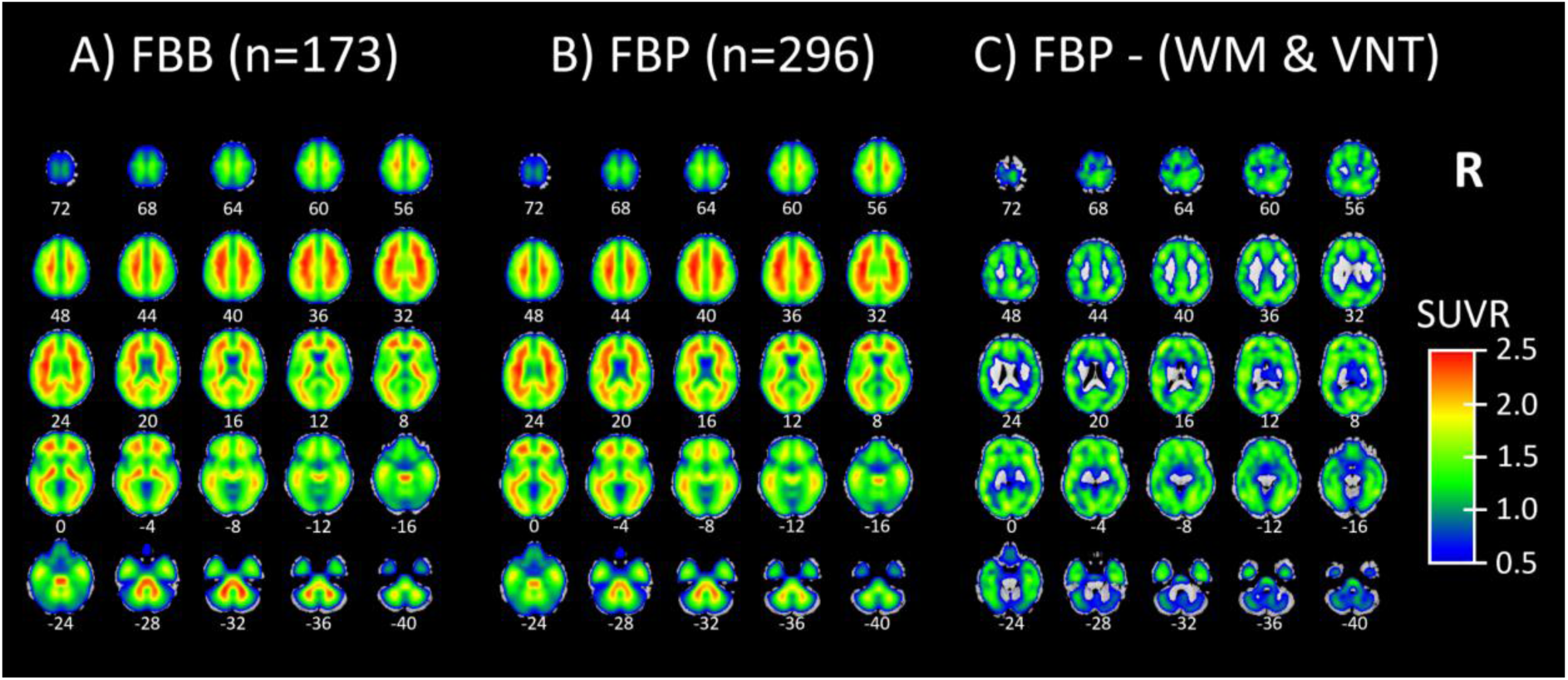
A) Molecular Image Brain Atlas (MIBA) average from 173 florbetaben participants. B) MIBA average of 296 florbetapir participants. C) Florbetapir MIBA average subtracted by white matter (WM) and ventricular (VN) related ICs (WM1, WM2, VN1, VN2, VN3, VN4 from Figure 2). In (C) it is seen that the subtraction of WM and ventricles leaves the SUVR signal in gray matter.

#### Regression of age with tracer uptake

According to previous literature (Fleisher et al., 2013, Bilgel et al., 2016, Jack et al., 2015) amyloids increase with age in gray matter, likely depicting specific binding to amyloid-beta plaques. Another complication for FBB and FBP is that the binding in WM may be related with beta-sheet structure of the myelin, decreasing the SUVR values (in white matter voxels) with subject age due to WM degradation (Catafau & Bullich, 2015; Moscoso et al., 2022). In addition, ventricle size may increase with age in older adults even for control participants (Padhy et al., 2014). This study targets the increased Aβ proteins, which increases with age in gray matter, but not white matter degradation that may decrease SUVR in FBP with age nor increased ventricular size which may decrease SUVR in regions close to CSF and ventricles in both FBB and FBP.

In this study all subjects DKT ROI’s SUVR and AβNs loading values were regressed with the subjects’ ages, yielding a standardized β and p-value for the DKT ROI’s SUVR and AβNs. The standardized β value and the false discovery rate (FDR) corrected p-value for each AβN regression are found in Table 2 and for the DKT ROI regressions in Table S1. The results shown in Table S1 show that binding in ventricles/CSF was negatively associated with age for both FBB and FBP. In addition, in FBP, but not in FBB, the WM had a negative association between age and SUVR. Even if WM does not have a negative age-association for FBB, there may still be a strong uptake, seen in Figure 3A, and this WM uptake may be unspecific to Aβ. Therefore, regions that decrease SUVR with age may be related with white matter degradation or enlarged ventricles suggesting potential off-target interactions or non-specific binding. Due to this off-target interactions and confounding nature, both DKR-ROIs and AβNs with negative association between age and SUVR or loading values are excluded from further quantitative analyses. The negative age-associations exclusions are seen as x-neg in the q-columns in Table 2 and in the “q FBP”-column in Table S1. This includes the AβNs exclusions of CC3 and SM3 for FBB and AU1, SM1, SM3, VIS3, DM2 and DM4 for FBP. In summary for both the FBB and FBP DKT ROIs, eleven non-gray matter regions were excluded (skull, air cavities, CSF, and white matter, labeled as “x-non-gm” in Table S1) and two reference regions were excluded (labeled as “reference” in Table S1). In addition, for the FBP tracer, eleven regions were excluded due to their negative association with age (labeled as “x-neg” in Table S1), which possibly indicates WM or CSF is bleeding into the GM ROI (i.e. partial volume effects) or perhaps another non-specific binding effect.

In this study we associated the average SUVR or loading values with participant ages. To evaluate the anatomical association with age using the DKT atlas, we performed a simple linear regression between the average SUVR within each DKT ROI and the participant age. Analogously, for the NeuroMark AβNs, we regressed the AβNs’ loading values against age. Since we use the simple linear regression for all regressions with age, all standardized betas are identical with the Pearson correlation. Since the ages are the same for the DKT ROIs and the AβNs, the corresponding standardized beta values are the same as correlations, which are unbiased between the DKT ROIs (Table S1) and the AβNs (Table 2). In addition, the p-values from the simple linear regression are identical with Pearsons correlation p-values as well in this case.

Here we highlight the most significant age association for the DKT ROI for the FBB tracer, which is in the right inferior temporal cortex (Figure 4, depicted in blue, R^2^=0.11, p < 0.001, uncorrected). Similarly, for the NeuroMark AβNs, we regressed the AβNs’ loading values against age. The strongest component association was observed in the CC1 AβN, which has its peak in the medial temporal cortex (Figure 4, depicted in orange, R^2^=0.18, p < 0.001, uncorrected).

**Figure 4:**
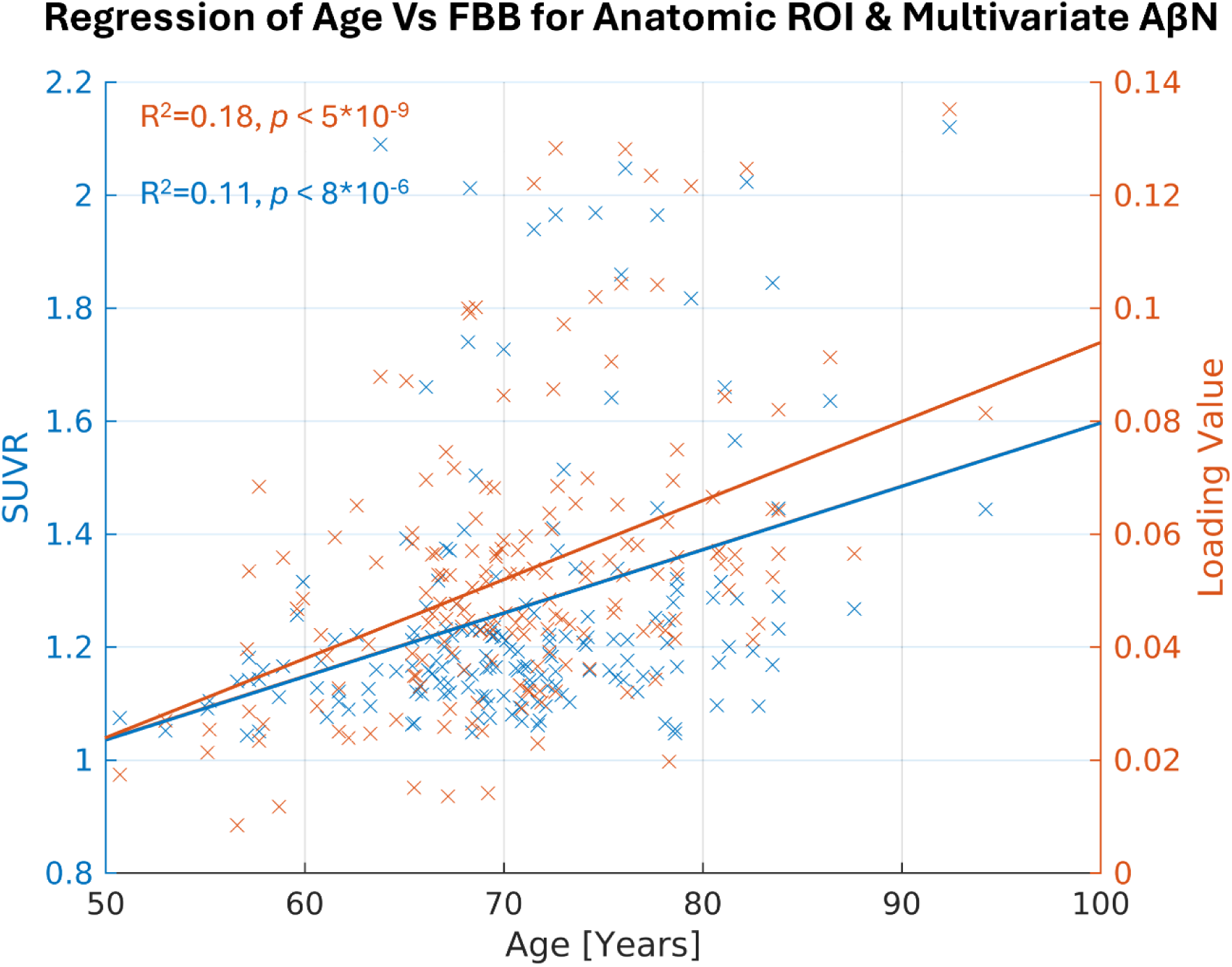
The most significant age association for anatomical ROI in blue and multivariate AβN in orange, making biological references for the atlas and the template. The anatomical ROI with the highest R^2^ (0.11) is the right inferior temporal cortex in the Desikan-Killiany-Tourville (DKT) atlas. The highest R^2^ (0.18) for the multivariate AβN is the CC1 in the NeuroMark FBP template.

In the remaining part of the results we explore the age association significance for both AβNs and DKT ROIs. In addition, we used the FDR corrected p-values (q-values) and sort the AβNs and DKT ROIs by age association significance (Figure 5). Because the number of tests is considerably lower for NeuroMark AβNs compared to DKT ROIs, the FDR adjustment might be less strict for AβNs, thereby aiding the detection of significant outcomes. As a result, this facilitates the comparison of binding associations with age for either ROIs or AβNs. In Figure 5, it is seen that the age association trajectories are different between the AβNs and the DKT ROIs. The AβNs are decomposed into fewer parts than the DKT ROIs (15 AβNs < 87 ROIs). Interestingly, the age association significance is distributed so that for the ROIs, more than 88% (77 of 87 ROIs) have significant age association (q < 0.05). Conversely, less than 67% (10 of 15) of the AβNs had significant association with age. However, it seems as the top age association AβNs contenders increase a lot in effect and may even be more significant than the top contending DKT ROIs.

**Figure 5:**
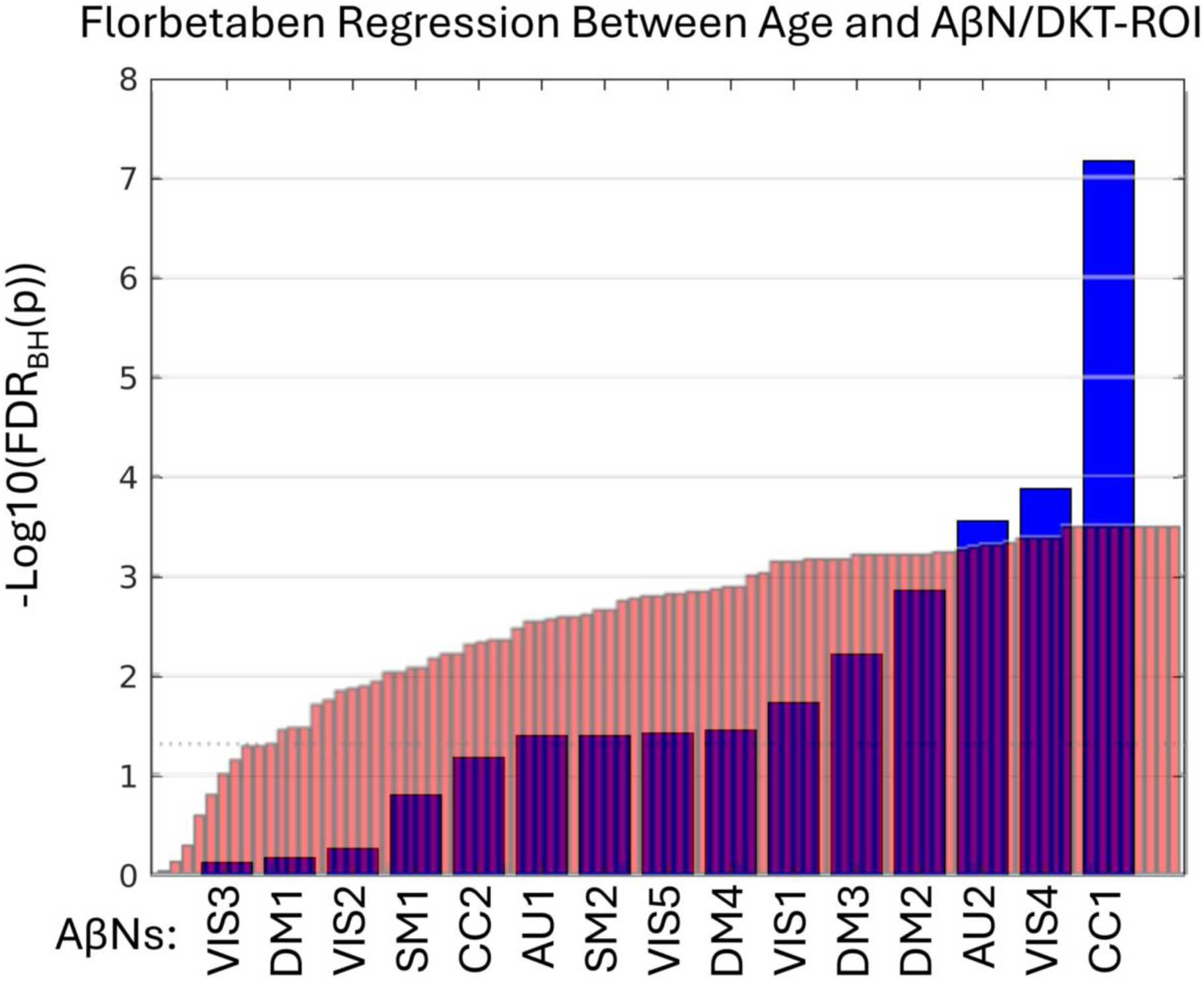
Y-axis show log10 of the FDR corrected p-values (q-value) after regressing 173 participants’ ages with FBB intensities from either amyloid-beta networks (AβNs, blue) or Desikan-Killiany-Tourville (DKT) ROIs (red). Only AβNs/DKT ROIs with positive age correlation and non-reference regions are included from the NeuroMark template AβNs or DKT ROIs from the FBB data. More stats are available in Tables 2 and S1. The dotted line at y = 1.3 represents a q = 0.05.

Since we used the 296 FBP participants to create the AβNs in our NeuroMark template, even as they are unsupervised and blind, we were careful to use its resulting loading values, but are hereafter interested in confirming that the components are capturing similar age association patterns as the FBB results. In addition, we explored if the relationship between age and AβN loading was consistent across these different age groups and tracer types (76.0 years for FBP and 70.8 years for FBB). All the q-values are ranked from lowest to highest and as seen in Figure 6. The CC1 component was the most significant for both FBP and FBB. However, it was slightly more significant for FBB, even though the NeuroMark AβNs were created from the FBP dataset. In addition, an AβNs pattern may have been identified, suggesting consistency across different PET tracers, since out of the 5 significant AβNs in FBP at least 4 were also significant in FBB (CC1, VIS4, VIS1, and SM2).

**Figure 6:**
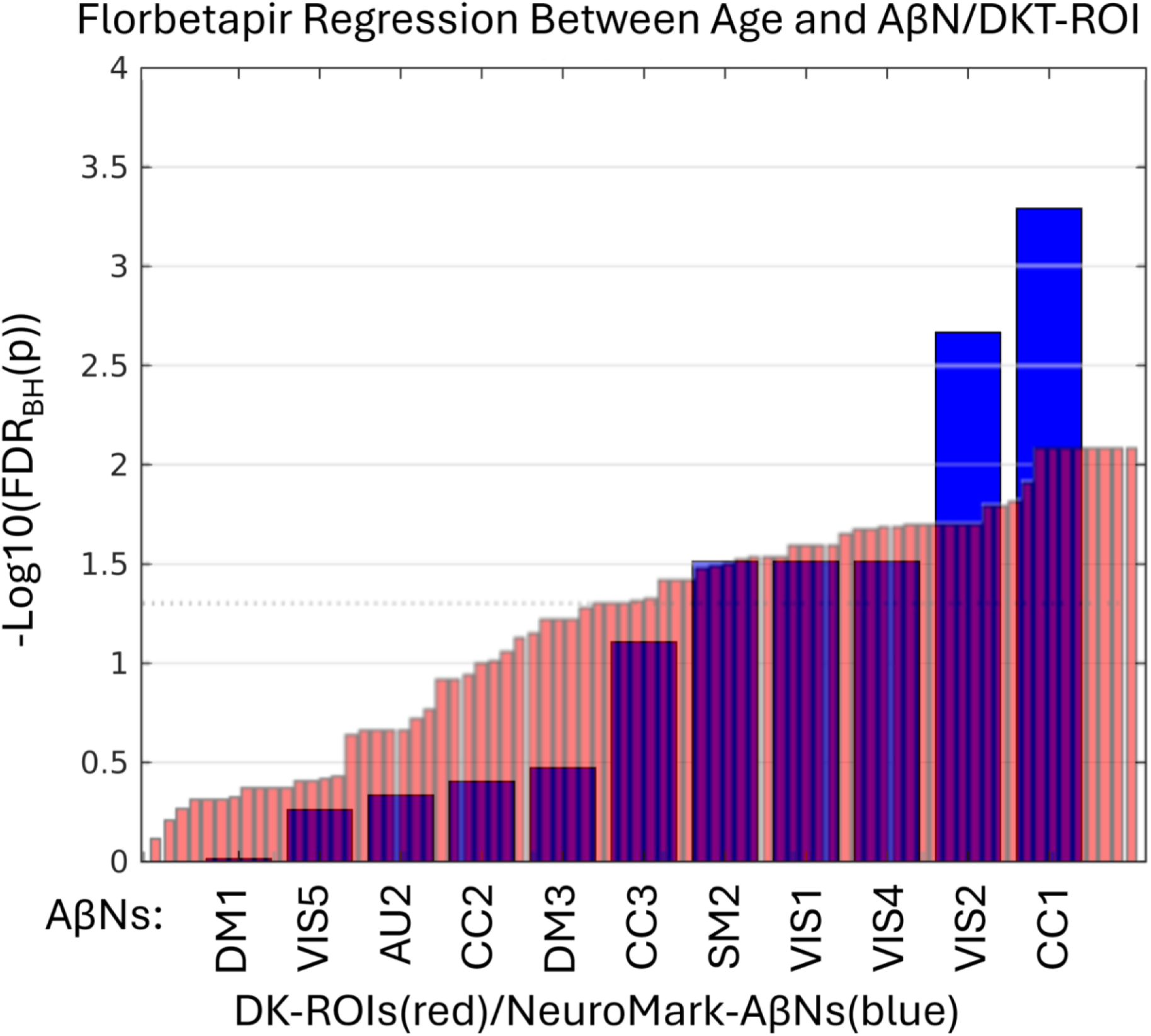
Y axis show log10 of the FDR corrected p-values (q-values) after regressing 296 participants’ ages with FBP intensities from either amyloid-beta networks (AβNs, blue) or Desikan-Killiany-Tourville (DKT) ROIs (red). Only AβNs/DKT ROIs with positive age correlation and non-reference regions are included from the NeuroMark template AβNs or DKT ROIs from the FBP data. More stats are available in Tables 2 and S1. The dotted line at y = 1.3 represents a q = 0.05.

## Discussion

In this study, we introduce NeuroMark PET, a novel method for computing replicable PET IC templates specifically designed for the FBP tracer. Utilizing the NeuroMark framework, we successfully extended our approach to the FBB tracer, which also targets Aβ pathology, thereby demonstrating the framework’s versatility across different tracers. Our findings indicate that NeuroMark PET can serve as a robust template for subsequent Aβ imaging studies. Additionally, we compared AβNs derived from our method to traditional anatomical atlases. Notably, the AβNs were decomposed into fewer components than anatomical ROIs, suggesting a more streamlined representation of amyloid distribution. Furthermore, the results indicate that AβNs may offer distinct advantages in capturing nuanced amyloid distribution patterns, potentially enhancing the sensitivity and specificity of Aβ detection.

### Previous Literature about the NeuroMark Florbetapir Amyloid β Networks and the Desikan Killiany Tourville ROIs

To get an impression of where the AβNs are mostly weighted, each AβN’s peak voxel was mapped to the DKT atlas, finding the most relevant DKT ROI for that AβN. Further on, up to two more DKT ROIs that were strongly weighted for each AβN were added, estimated qualitatively, characterizing each AβN anatomically. These anatomical DKT characteristics of the AβN are listed in Table S2. The identified DKT ROIs have been referenced in the literature for being related with mental impairments and various functions. I.e., for the CC1 AβN, which is strongly weighted in the inferior temporal cortex (IT, Table S2), potentially has the highest local amyloid-beta/tau interactions and a connectivity profile conducive to accelerating tau propagation (Lee et al., 2022). Additionally, the IT shows significant increases in Alzheimer’s patients when scanned with AV1451 targeting tau (Johnson et al., 2016) and with florbetaben targeting amyloids (Cho et al., 2016). Interestingly, the DKT ROI with the highest age effect was the right inferior temporal cortex. It may be noted that the NeuroMark AβNs, were all bilateral except from VIS1 that was dominant in the right hemisphere and VIS2 that was dominant in the left hemisphere. In addition, Pereira et al. (2020), using the PET Tau radioligand, found three unilateral out of ten independent components and two of them were very similar with the VIS1 and VIS2. Other components were similar between Pereira et al. (2020) and our study as well. A more comprehensive overview elucidating the primary anatomical locations and their functional roles within the DKT framework, offering an improved understanding of how AβNs interact with specific DKT ROIs is found in the supplementary Table S2 and under section “Supplementary Literature of NeuroMark Amyloid β Networks & DKT Anatomy.”

### Automation and Advantages of the NeuroMark PET Pipeline

The NeuroMark PET Pipeline can effectively eliminate the need for model order selection and component identification procedures, while allowing for data-driven approaches to retain more individualized variability. By utilizing a NeuroMark template as a spatial prior, we ensure the correspondence of extracted functional network features across diverse analyses, studies, and datasets, which will definitely facilitate the generation of reproducible findings that might advance the reliability and robustness in neuroimaging research.

Given that NeuroMark is a robust and fully automated framework that continuously offers reliable templates for multiple modalities, unexplored multimodal areas could present valuable opportunities for future research. The NeuroMark FBP template is a novel development as it establishes a reliable foundation extended to be used for PET modalities, facilitating multimodal analyses in an automated fashion. The ICA approach employed in NeuroMark enables the identification of independent networks with co-varied voxels across participants, and also helps to separate out artifacts and partial volume effects. Notably, traditional ROIs-based approaches relied on predefined but fixed brain regions across subjects and scans which might therefore neglect the variation in the alignments of ROIs with the underlying chemo-architectural boundaries. In contrast, the data-driven NeuroMark approach offers a flexible alternative that accurately reflects the intrinsic brain architecture according to individual scan, making extracted PET outputs more reliable and accessible to non-experts, such as researchers from other fields, including clinicians.

### Independent Component Analysis Empirical Spatial Findings

As mentioned above, the NeuroMark framework measures AβNs reproducibility (in this study by measuring paired AβNs spatial correlation across groups A and B). The most reproducible component was the first white matter AβN (Figure 7), however this component was excluded because it is not considered relevant to the research question (having no correspondence to gray matter). The FBP radioligands have substantial non-specific binding in the white matter. In agreement with previous literature about myelin integrity (Catafau & Bullich, 2015) we found a negative association with age for this WM IC. That the white matter is clearly detected and delineated provides evidence that ICs may correspond to biologically distinct structures, implying that the ICA is detecting relevant patterns in the underlying signal. Analogously several biologically relevant ventricle components were detected by ICA as well.

**Figure 7.**
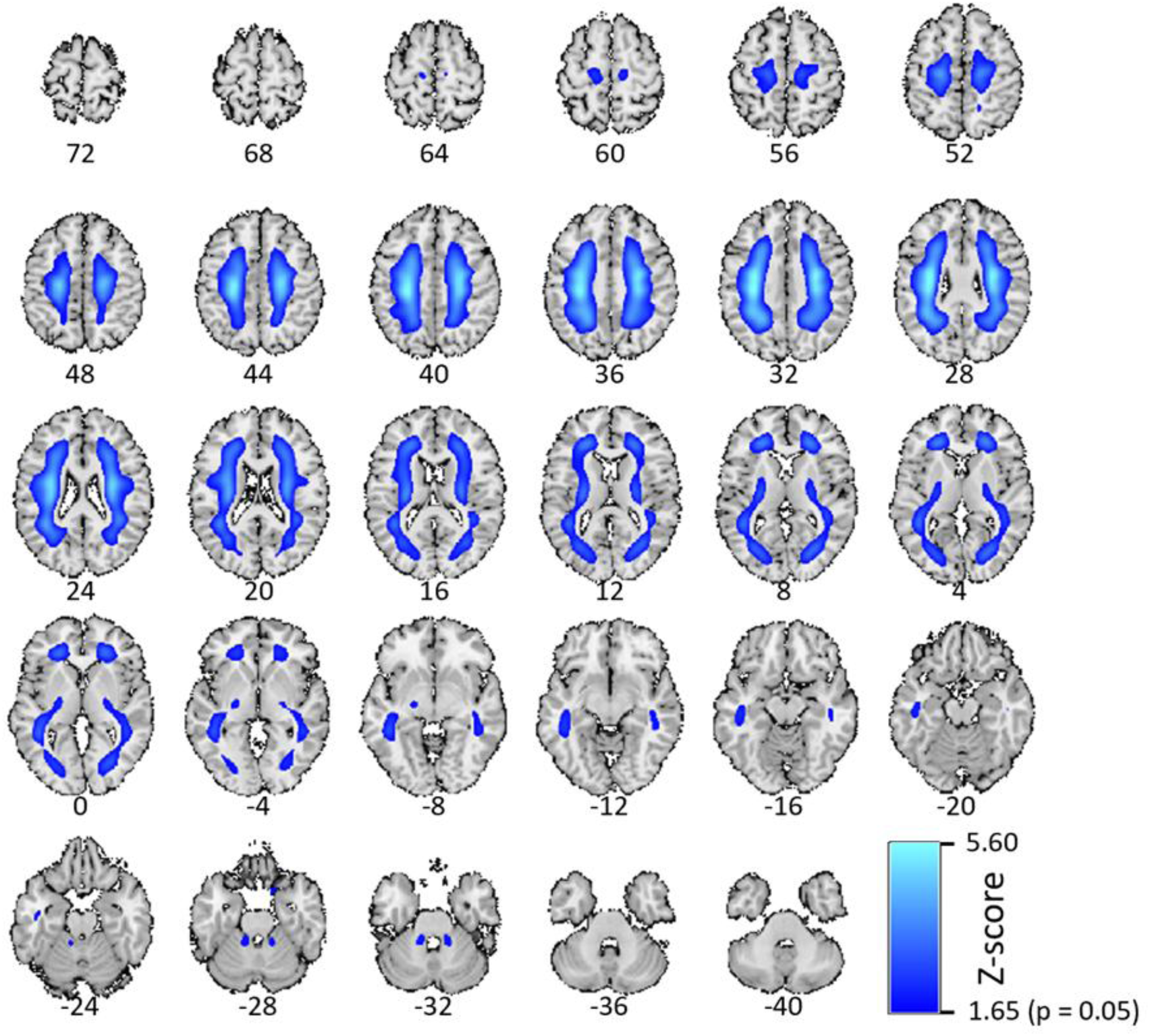
A white matter component correlated best between groups A and B and may be the most reproducible ICA component for the florbetapir radioligand.

### Limitations

Using different tracers, we found many similarities such that the CC1 was the most significant AβN is both FBB and FBP. However, a difference between the tracers was that they had slightly different negative age associations. For FBB the CC3 and SM3 had negative age effects, while for FBP, DM2, DM4, AU1, SM1, and VIS3 had negative age effects. The negative ICs were excluded from this study. The different patterns of negative age effects may be related to a slightly different molecular binding. E.g., FBP had a negative association between age and SUVR in WM, but not FBB (also seen in Table S1).

In this study we compare NeuroMark AβNs to DKT anatomical ROIs. To calculate partial volume correction (PVC) for DKT ROIs one has to use a compartment-based method and for NeuroMark AβNs one has to calculate PVC using a voxel-based method. To make sure we treated the processing the same for the anatomical atlas and for the NeuroMark template, we decided not to use any PVC in this study. However, PVC is an important factor and we may explore how PVC affects the results in a future study.

Another difference is how the DKT ROIs and the AβNs represent features. It may be posited that the larger AβNs offer enhanced stability relative to the smaller DKT ROIs, potentially rendering the latter noisier, it is essential to emphasize the role of ICA as an effective dimension reduction tool. ICA excels in simplifying complex datasets by decomposing them into a limited number of meaningful components, thereby enhancing interpretability and reducing noise. This dimensionality reduction not only benefits researchers by streamlining data analysis but may also preserve the integrity of significant neural patterns. Comparing the 15 NeuroMark AβNs, which are considerably larger in size to the 87 statistically significant DKT ROIs, the CC1 AβN exhibited greater significance than the right inferior temporal cortex ROI, as illustrated in Figure 4. The overlapping and weighted nature of ICA-derived components provides a complementary perspective to traditional ROI-based approaches. An approach that adheres to established scientific standards and offers researchers an additional angle to uncover intricate neural relationships.

## Conclusions

This study introduces and validates the NeuroMark PET approach, leveraging spatially constrained independent component analysis to generate biologically meaningful and reproducible brain networks. By applying this method, we demonstrated that NeuroMark PET may complement the traditional atlas-based analysis (DKT). We demonstrate by showing an example of capturing age-related binding changes in the brain, specifically in cortical regions relevant to neurodegenerative processes. NeuroMark PET can at least have three advantages to an anatomical atlas (e.g., DKT). 1) A few NeuroMark AβNs may excel as most significant compared to the most significant DKT ROIs, which could be crucial for applications where detecting leading signals is more valuable than a broad but less intense detection. 2) The complexity of the measured Aβ uptake, across subject populations, may be decomposed, by NeuroMark ICA, into a limited number of contrasting AβNs, each aligning to an Aβ uptake subpattern representing an important aspect. This automated ICA approach allows for overlapping regions, reducing the complexity and potential bias associated with manual data division. 3) The NeuroMark AβNs are spatial patterns customized specifically for Aβ uptake. In addition, NeuroMark can customize maps for other radioligands as well. Conversely, the anatomical atlases are rigid, having the same fixed borders for every radioligand.

Strengths above are likely related to the fact that ICA is data-driven, capturing voxels that show covariation among participants in maximally spatially independent maps. The ICA results also benefit from data-reduction, as the multiple comparison burden is lower for ICA compared to the ROI analysis.

The NeuroMark AβN template, which excludes non gray matter regions (e.g., white matter and ventricles), may therefore provide a more efficient and biologically relevant representation of molecular tracer distributions. This advancement highlights the potential of NeuroMark AβNs to enhance our understanding of the molecular underpinnings of brain function and pathology, paving the way for more sensitive and individualized analyses in large-scale multimodal datasets. This approach offers a straightforward and objective way to generate radioligand specific maps alternative to conventional region of interest (ROI)-based methodologies, emphasizing the importance of data-driven techniques in neuroimaging research.

## Data availability

The data that supports the findings of this study are available on request from the public Alzheimer’s Disease Neuroimaging Initiative (ADNI). Application to access the ADNI dataset is found at https://adni.loni.usc.edu/data-samples/adni-data/#AccessData.

## Ethics approval statement

This study utilized data from the ADNI database. ADNI has obtained all necessary IRB approvals and met all ethical standards in the collection of their data. As per ADNI protocols, all procedures performed in studies involving human participants were in accordance with the ethical standards of the institutional and/or national research committee and with the 1964 Helsinki declaration and its later amendments or comparable ethical standards. The use of ADNI data in this study was approved through an online application process, and all investigators agreed to the ADNI Data Use Agreement, which includes provisions for data confidentiality and ethical use.

## Patient consent and permission to reproduce material from ADNI

Data used in the preparation of this study were obtained from the ADNI database. The investigators within ADNI obtained written informed consent from all participants, and approval was granted by the relevant institutional review boards. The authors acknowledge permission to use and reproduce ADNI data in accordance with ADNI’s data use policies. In addition, our study was approved by the ADNI Data and Publications Committee (DPC).

## Additional acknowledgements

Data collection and sharing for this project were funded by the ADNI (National Institutes of Health Grant U01 AG024904) and DOD ADNI (Department of Defense award number W81XWH-12-2-0012). ADNI is funded by the National Institute on Aging, the National Institute of Biomedical Imaging and Bioengineering, and through generous contributions from the following: AbbVie, Alzheimer’s Association; Alzheimer’s Drug Discovery Foundation; Araclon Biotech; BioClinica, Inc.; Biogen; Bristol-Myers Squibb Company; CereSpir, Inc.; Cogstate; Eisai Inc.; Elan Pharmaceuticals, Inc.; Eli Lilly and Company; EuroImmun; F. Hoffmann-La Roche Ltd and its affiliated company Genentech, Inc.; Fujirebio; GE Healthcare; IXICO Ltd.; Janssen Alzheimer Immunotherapy Research & Development, LLC.; Johnson & Johnson Pharmaceutical Research & Development LLC.; Lumosity; Lundbeck; Merck & Co., Inc.; Meso Scale Diagnostics, LLC.; NeuroRx Research; Neurotrack Technologies; Novartis Pharmaceuticals Corporation; Pfizer Inc.; Piramal Imaging; Servier; Takeda Pharmaceutical Company; and Transition Therapeutics. The Canadian Institutes of Health Research is providing funds to support ADNI clinical sites in Canada. Private sector contributions are facilitated by the Foundation for the National Institutes of Health (www.fnih.org). The grantee organization is the Northern California Institute for Research and Education, and the study is coordinated by the Alzheimer’s Therapeutic Research Institute at the University of Southern California. ADNI data are disseminated by the Laboratory for Neuro Imaging at the University of Southern California.

