## Supplemental Methods for "Building Multivariate Molecular Imaging Brain Atlases Using the NeuroMark PET Independent Component Analysis Framework"

### Supplementary Material

#### Supplementary Methods

ADNI search criteria for fMRI and PET images was as follows. As of September 28, 2023, 438 cognitively normal ADNI participants underwent FBP PET scans, and the radiopharmaceutical 18F-AV45. Simultaneously, 4,826  T1-weighted MRI scans were found for the FBP participants with the following criteria: T1 weighting, field strength between 1.4 and 3.1 Tesla, and slice thickness between 0.9 and 1.3 mm. As of December 11, 2023, 214 participant FBB PET scans were identified as cognitive normal using the analogous search criteria as FBP with the exception of the radiopharmaceutical parameter, which was selected to only 18F-FBB. 537 T1-weighted MRI scans were found with the analogous criteria as FBP, but now for the FBB participants. All resulting PET (FBP and FBB) images had 4 frames with varying slice thicknesses between 1.0 mm and 3.4 mm and all the downloaded T1w MRI were scanned with a slice thickness of either 1 mm or 1.2mm at a 1.5 or 3 Tesla scanner. Matching the PET and T1w results for paired processing, resulted in 322 cognitive normal FBP and T1w MRIs being properly matched for processing and 198 cognitive normal FBB images were matched with T1w MRIs as well.
