## Supplemental Table S1 for "Building Multivariate Molecular Imaging Brain Atlases Using the NeuroMark PET Independent Component Analysis Framework"

### Supplementary Material

**Table S1**. Descriptive statistics for the SUVR values in the DKT ROIs: mean (μ), standard deviation (σ), association with age using beta (β) and the FDR corrected p (q) for FBB and FBP. In the q columns some DKT ROIs are excluded: because they are not gray matter (x-non-gm), because they were used for SUVR reference or because the age association was negative (x-neg). Acronyms: left (L.); right (R.).

| DKT Label | μ FBB | σ FBB | β FBB | q FBB | μ FBP | σ FBP | β FBP | q FBP |
|---|---|---|---|---|---|---|---|---|
| L. Cerebral White Matter | 2.00 | 0.17 | 0.114 | x-non-gm | 2.02 | 0.23 | -0.118 | x-non-gm |
| L. Cerebellum White Matter | 2.02 | 0.16 | 0.163 | x-non-gm | 1.90 | 0.20 | -0.167 | x-non-gm |
| L. Cerebellum Cortex | 1.00 | 0.01 | 0.067 | reference | 1.00 | 0.01 | -0.206 | reference |
| L. Thalamus | 1.52 | 0.16 | 0.127 | 1.00E-01 | 1.44 | 0.16 | -0.087 | x-neg |
| L. Caudate | 1.18 | 0.15 | 0.047 | 5.50E-01 | 1.23 | 0.17 | 0.056 | 3.91E-01 |
| L. Putamen | 1.48 | 0.18 | 0.274 | 7.08E-04 | 1.51 | 0.20 | 0.130 | 4.88E-02 |
| L. Pallidum | 1.93 | 0.19 | 0.206 | 8.66E-03 | 1.82 | 0.21 | -0.133 | x-neg |
| Brain Stem | 1.72 | 0.14 | 0.224 | 4.47E-03 | 1.60 | 0.16 | -0.081 | x-neg |
| L. Hippocampus | 1.26 | 0.10 | 0.001 | 9.85E-01 | 1.23 | 0.12 | -0.320 | x-neg |
| L. Amygdala | 1.16 | 0.11 | 0.190 | 1.51E-02 | 1.17 | 0.14 | 0.037 | 5.41E-01 |
| CSF | 0.85 | 0.16 | -0.370 | x-non-gm | 0.82 | 0.17 | -0.441 | x-non-gm |
| L. Accumbens area | 1.21 | 0.27 | 0.162 | 3.73E-02 | 1.25 | 0.28 | 0.159 | 2.12E-02 |
| L. VentralDC | 1.82 | 0.16 | 0.183 | 1.88E-02 | 1.70 | 0.18 | -0.17 | x-neg |
| L. choroid plexus | 1.04 | 0.19 | -0.287 | x-non-gm | 0.97 | 0.19 | -0.387 | x-non-gm |
| R. Cerebral White Matter | 2.02 | 0.18 | 0.147 | x-non-gm | 2.03 | 0.23 | -0.111 | x-non-gm |
| R. Cerebellum White Matter | 2.05 | 0.16 | 0.14 | x-non-gm | 1.92 | 0.20 | -0.152 | x-non-gm |
| R. Cerebellum Cortex | 1.00 | 0.01 | -0.066 | reference | 1.00 | 0.01 | 0.208 | reference |
| R. Thalamus | 1.52 | 0.16 | 0.15 | 5.37E-02 | 1.44 | 0.16 | -0.074 | x-neg |
| R. Caudate | 1.20 | 0.16 | 0.085 | 2.74E-01 | 1.25 | 0.17 | 0.043 | 4.93E-01 |
| R. Putamen | 1.48 | 0.19 | 0.332 | 3.21E-04 | 1.51 | 0.19 | 0.137 | 3.82E-02 |
| R. Pallidum | 1.93 | 0.19 | 0.239 | 2.62E-03 | 1.82 | 0.21 | -0.135 | x-neg |
| R. Hippocampus | 1.28 | 0.09 | 0.02 | 8.04E-01 | 1.25 | 0.12 | -0.344 | x-neg |
| R. Amygdala | 1.19 | 0.12 | 0.223 | 4.70E-03 | 1.18 | 0.14 | 0.016 | 7.80E-01 |
| R. Accumbens area | 1.23 | 0.27 | 0.235 | 2.93E-03 | 1.27 | 0.27 | 0.145 | 3.02E-02 |
| R. VentralDC | 1.79 | 0.16 | 0.15 | 5.34E-02 | 1.68 | 0.18 | -0.187 | x-neg |
| R. choroid plexus | 1.07 | 0.17 | -0.272 | x-non-gm | 1.02 | 0.17 | -0.365 | x-non-gm |
| AirCavity | 0.50 | 0.10 | 0.123 | x-non-gm | 0.53 | 0.12 | 0.069 | x-non-gm |
| Skull | 0.82 | 0.18 | -0.07 | x-non-gm | 0.74 | 0.18 | -0.092 | x-non-gm |
| Vermis | 0.95 | 0.04 | 0.107 | 1.69E-01 | 0.95 | 0.05 | -0.108 | x-neg |

| | | | | | | | |
|---|---|---|---|---|---|---|---|
| Pons | 2.17 | 0.19 | 0.256 | 1.45E-03 | 2.04 | 0.24 | -0.109 | x-neg |
| CSF Extra Cerebral | 0.89 | 0.15 | 0.102 | x-non-gm | 0.88 | 0.15 | 0.059 | x-non-gm |
| Head Extra Cerebral | 1.02 | 0.15 | -0.106 | x-non-gm | 1.04 | 0.18 | -0.118 | x-non-gm |
| L. Banks of Sup. Temp. Sulcus | 1.47 | 0.28 | 0.258 | 1.36E-03 | 1.57 | 0.32 | 0.137 | 3.82E-02 |
| L. Caudal Anterior Cingulate | 1.39 | 0.25 | 0.298 | 4.30E-04 | 1.44 | 0.26 | 0.147 | 2.90E-02 |
| L. Caudal Middle Frontal | 1.34 | 0.24 | 0.259 | 1.31E-03 | 1.40 | 0.28 | 0.2 | 8.22E-03 |
| L. Cuneus | 1.29 | 0.16 | 0.205 | 8.81E-03 | 1.33 | 0.21 | 0.046 | 4.72E-01 |
| L. Entorhinal Cortex | 1.10 | 0.13 | 0.211 | 7.13E-03 | 1.07 | 0.13 | 0.058 | 3.86E-01 |
| L. Fusiform Gyrus | 1.26 | 0.20 | 0.251 | 1.65E-03 | 1.31 | 0.24 | 0.128 | 5.03E-02 |
| L. Inferior Parietal Lobule | 1.34 | 0.26 | 0.267 | 9.41E-04 | 1.40 | 0.29 | 0.131 | 4.78E-02 |
| L. Inferior Temporal Gyrus | 1.28 | 0.24 | 0.280 | 6.18E-04 | 1.33 | 0.28 | 0.168 | 2.01E-02 |
| L. Isthmus of Cingulate Gyrus | 1.40 | 0.25 | 0.322 | 3.21E-04 | 1.42 | 0.26 | 0.143 | 3.23E-02 |
| L. Lateral Occipital Cortex | 1.31 | 0.17 | 0.282 | 6.18E-04 | 1.36 | 0.25 | 0.081 | 2.17E-01 |
| L. Lateral Orbitofrontal Cortex | 1.33 | 0.24 | 0.274 | 7.14E-04 | 1.37 | 0.26 | 0.184 | 1.25E-02 |
| L. Lingual Gyrus | 1.21 | 0.14 | 0.203 | 9.38E-03 | 1.25 | 0.21 | 0.052 | 4.27E-01 |
| L. Medial Orbitofrontal Cortex | 1.27 | 0.27 | 0.251 | 1.65E-03 | 1.31 | 0.28 | 0.193 | 8.22E-03 |
| L. Middle Temporal Gyrus | 1.25 | 0.24 | 0.274 | 7.08E-04 | 1.29 | 0.27 | 0.152 | 2.55E-02 |
| L. Parahippocampal Gyrus | 1.15 | 0.17 | 0.237 | 2.73E-03 | 1.15 | 0.18 | 0.051 | 4.28E-01 |
| L. Paracentral Lobule | 1.34 | 0.20 | 0.166 | 3.41E-02 | 1.37 | 0.23 | 0.077 | 2.29E-01 |
| L. Pars Opercularis | 1.29 | 0.24 | 0.255 | 1.47E-03 | 1.35 | 0.26 | 0.157 | 2.22E-02 |
| L. Pars Orbitalis | 1.28 | 0.25 | 0.253 | 1.56E-03 | 1.29 | 0.27 | 0.169 | 2.01E-02 |
| L. Pars Triangularis | 1.34 | 0.24 | 0.248 | 1.83E-03 | 1.38 | 0.27 | 0.147 | 2.90E-02 |
| L. Pericalcarine Cortex | 1.38 | 0.18 | 0.181 | 2.04E-02 | 1.48 | 0.26 | 0.108 | 9.79E-02 |
| L. Postcentral Gyrus | 1.24 | 0.18 | 0.139 | 7.37E-02 | 1.27 | 0.20 | 0.057 | 3.91E-01 |
| L. Posterior Cingulate | 1.41 | 0.28 | 0.288 | 5.68E-04 | 1.46 | 0.30 | 0.153 | 2.55E-02 |
| L. Precentral Gyrus | 1.31 | 0.16 | 0.197 | 1.19E-02 | 1.34 | 0.19 | 0.090 | 1.71E-01 |
| L. Precuneus | 1.37 | 0.31 | 0.265 | 1.01E-03 | 1.44 | 0.33 | 0.138 | 3.82E-02 |
| L. Rostral Anterior Cingulate | 1.30 | 0.28 | 0.280 | 6.18E-04 | 1.36 | 0.30 | 0.166 | 2.01E-02 |
| L. Rostral Middle Frontal | 1.32 | 0.29 | 0.243 | 2.22E-03 | 1.37 | 0.31 | 0.175 | 1.63E-02 |
| L. Superior Frontal Gyrus | 1.27 | 0.26 | 0.250 | 1.70E-03 | 1.32 | 0.27 | 0.198 | 8.22E-03 |
| L. Superior Parietal Lobule | 1.29 | 0.23 | 0.194 | 1.30E-02 | 1.35 | 0.26 | 0.107 | 9.93E-02 |
| L. Superior Temporal Gyrus | 1.24 | 0.21 | 0.238 | 2.62E-03 | 1.28 | 0.24 | 0.129 | 5.01E-02 |
| L. Supramarginal Gyrus | 1.30 | 0.25 | 0.224 | 4.47E-03 | 1.36 | 0.27 | 0.116 | 7.42E-02 |
| L. Frontal Pole | 1.13 | 0.30 | 0.253 | 1.56E-03 | 1.13 | 0.30 | 0.193 | 8.22E-03 |
| L. Temporal Pole | 1.14 | 0.16 | 0.230 | 3.49E-03 | 1.14 | 0.18 | 0.161 | 2.06E-02 |
| L. Transverse Temporal Gyrus | 1.28 | 0.21 | 0.153 | 5.03E-02 | 1.34 | 0.24 | 0.100 | 1.23E-01 |
| L. Insula | 1.24 | 0.20 | 0.235 | 2.95E-03 | 1.27 | 0.21 | 0.122 | 5.97E-02 |
| R. Banks of Sup. Temp. Sulcus | 1.48 | 0.28 | 0.298 | 4.30E-04 | 1.58 | 0.32 | 0.147 | 2.90E-02 |
| R. Caudal Anterior Cingulate | 1.36 | 0.26 | 0.275 | 7.08E-04 | 1.41 | 0.27 | 0.122 | 5.99E-02 |
| R. Caudal Middle Frontal | 1.34 | 0.25 | 0.275 | 7.08E-04 | 1.40 | 0.27 | 0.195 | 8.22E-03 |
| R. Cuneus | 1.28 | 0.16 | 0.242 | 2.30E-03 | 1.31 | 0.20 | 0.042 | 4.93E-01 |
| R. Entorhinal Cortex | 1.10 | 0.13 | 0.313 | 3.21E-04 | 1.07 | 0.13 | 0.030 | 6.19E-01 |
| R. Fusiform Gyrus | 1.26 | 0.22 | 0.308 | 3.21E-04 | 1.29 | 0.23 | 0.123 | 5.97E-02 |
| R. Inferior Parietal Lobule | 1.34 | 0.28 | 0.282 | 6.18E-04 | 1.40 | 0.29 | 0.141 | 3.33E-02 |

| | | | | | | | | |
|---|---|---|---|---|---|---|---|---|
| R. Inferior Temporal Gyrus | 1.27 | 0.25 | 0.333 | 3.21E-04 | 1.32 | 0.26 | 0.175 | 1.63E-02 |
| R. Isthmus of Cingulate Gyrus | 1.40 | 0.26 | 0.310 | 3.21E-04 | 1.42 | 0.26 | 0.144 | 3.19E-02 |
| R. Lateral Occipital Cortex | 1.33 | 0.20 | 0.292 | 5.03E-04 | 1.35 | 0.24 | 0.100 | 1.23E-01 |
| R. Lateral Orbitofrontal Cortex | 1.34 | 0.25 | 0.311 | 3.21E-04 | 1.38 | 0.25 | 0.179 | 1.52E-02 |
| R. Lingual Gyrus | 1.22 | 0.17 | 0.215 | 6.32E-03 | 1.23 | 0.18 | 0.051 | 4.28E-01 |
| R. Medial Orbitofrontal Cortex | 1.28 | 0.29 | 0.295 | 4.70E-04 | 1.33 | 0.29 | 0.193 | 8.22E-03 |
| R. Middle Temporal Gyrus | 1.26 | 0.25 | 0.309 | 3.21E-04 | 1.31 | 0.26 | 0.161 | 2.06E-02 |
| R. Parahippocampal Gyrus | 1.17 | 0.18 | 0.310 | 3.21E-04 | 1.17 | 0.18 | 0.050 | 4.28E-01 |
| R. Paracentral Lobule | 1.34 | 0.20 | 0.193 | 1.36E-02 | 1.38 | 0.23 | 0.086 | 1.91E-01 |
| R. Pars Opercularis | 1.31 | 0.25 | 0.280 | 6.18E-04 | 1.36 | 0.26 | 0.164 | 2.01E-02 |
| R. Pars Orbitalis | 1.29 | 0.26 | 0.302 | 4.27E-04 | 1.30 | 0.27 | 0.165 | 2.01E-02 |
| R. Pars Triangularis | 1.35 | 0.26 | 0.273 | 7.14E-04 | 1.38 | 0.27 | 0.151 | 2.55E-02 |
| R. Pericalcarine Cortex | 1.37 | 0.20 | 0.215 | 6.32E-03 | 1.45 | 0.25 | 0.080 | 2.18E-01 |
| R. Postcentral Gyrus | 1.24 | 0.18 | 0.164 | 3.55E-02 | 1.27 | 0.20 | 0.041 | 4.97E-01 |
| R. Posterior Cingulate | 1.40 | 0.29 | 0.313 | 3.21E-04 | 1.46 | 0.29 | 0.153 | 2.55E-02 |
| R. Precentral Gyrus | 1.31 | 0.17 | 0.204 | 9.36E-03 | 1.35 | 0.19 | 0.080 | 2.18E-01 |
| R. Precuneus | 1.37 | 0.30 | 0.272 | 7.37E-04 | 1.43 | 0.32 | 0.126 | 5.33E-02 |
| R. Rostral Anterior Cingulate | 1.31 | 0.29 | 0.290 | 5.21E-04 | 1.38 | 0.29 | 0.159 | 2.12E-02 |
| R. Rostral Middle Frontal | 1.34 | 0.30 | 0.286 | 5.80E-04 | 1.38 | 0.31 | 0.167 | 2.01E-02 |
| R. Superior Frontal Gyrus | 1.28 | 0.26 | 0.279 | 6.37E-04 | 1.32 | 0.27 | 0.195 | 8.22E-03 |
| R. Superior Parietal Lobule | 1.29 | 0.22 | 0.240 | 2.56E-03 | 1.34 | 0.26 | 0.080 | 2.18E-01 |
| R. Superior Temporal Gyrus | 1.24 | 0.21 | 0.280 | 6.18E-04 | 1.27 | 0.23 | 0.128 | 5.07E-02 |
| R. Supramarginal Gyrus | 1.29 | 0.23 | 0.259 | 1.31E-03 | 1.35 | 0.26 | 0.111 | 8.84E-02 |
| R. Frontal Pole | 1.17 | 0.31 | 0.287 | 5.74E-04 | 1.17 | 0.30 | 0.212 | 8.22E-03 |
| R. Temporal Pole | 1.14 | 0.17 | 0.300 | 4.30E-04 | 1.13 | 0.17 | 0.103 | 1.14E-01 |
| R. Transverse Temporal Gyrus | 1.26 | 0.20 | 0.220 | 5.14E-03 | 1.31 | 0.22 | 0.060 | 3.77E-01 |
| R. Insula | 1.25 | 0.21 | 0.293 | 5.01E-04 | 1.26 | 0.21 | 0.117 | 7.11E-02 |
