## Supplemental Table S2 for "Building Multivariate Molecular Imaging Brain Atlases Using the NeuroMark PET Independent Component Analysis Framework"

### Supplementary Material

**Table S2**. Lists the each Amyloid β Network (AβN) and its peak related with the Desikan Killiani Tourville (DKT) atlas under the "AβN Peak Location" column. Then a couple of additional DKT ROIs that are strongly associated with the AβN may be listed under the "2nd Relevant DKT ROI" and "3rd Relevant DKT ROI" columns. Acronyms: left (L.); right (R.).

| AβN | AβN Peak Location | 2nd Relevant DKT ROI | 3rd Relevant DKT ROI |
|---|---|---|---|
| AU1 | Supramarginal Gyri | Postcentral Gyri | Insular Cortices |
| AU2 | Transverse Temporal Gyri | Superior Temporal Gyri | |
| SM1 | Superior Parietal Lobules | Inferior Parietal Lobules | |
| SM2 | Precentral Gyri | Superior Frontal Gyri | Caudal Middle Frontal Cortices |
| SM3 | Postcentral Gyri | Superior Parietal Lobules | Paracentral Lobules |
| VIS1 | L. Lateral Occipital Cortex | L. Lingual Gyrus | L. Pericalcarine Cortex |
| VIS2 | R. Lateral Occipital Cortex | | |
| VIS3 | Lateral Occipital Cortices | Superior Parietal Lobules | Cuneus Cortices |
| VIS4 | Fusiform Gyri | Inferior Temporal Gyri | Lateral Occipital Cortices |
| VIS5 | Pericalcarine Cortices | Cuneus Cortices | Lingual Gyri |
| CC1 | Medial Temporal Gyri | Inferior Temporal Gyri | Superior Temporal Gyri |
| CC2 | Rostral Mid. Frontal Cortices | Superior Frontal Gyri | Frontal Pole Cortices |
| CC3 | Superior Frontal Gyri | Precentral Gyri | |
| DM1 | Paracentral Lobules | Superior Frontal Gyri | |
| DM2 | Isthmus of Cingulate | Precuneus Cortices | |
| DM3 | Precuneus Cortices | | |
| DM4 | Rostral Ant. Cingulate Cortices | Caudal Ant. Cingulate Cortices | Superior Frontal Gyri |

#### Supplementary Literature of NeuroMark Amyloid β Networks & DKT Anatomy

This section offers an overview of the NeuroMark AβNs, emphasizing their peak intensities relative to DKT ROIs and a couple additional relevant DKT ROIs listed in Table S2. All components are bilateral unless specified. The DKT ROIs are ordered by the extent of coverage by each AβN, starting with the ROI containing the AβN peak.

The CC1 AβN have peaks located in the middle temporal gyri (MT), strongly covering the inferior temporal gyri (IT) and the superior temporal gyri (ST). The MT and IT play a crucial role in semantic verbal memory processing (Bartha et al., 2003). More information about IT is found under the Discussion section. The IT are visual pathways implicated in object, face, and scene perception (Conway et al., 2018). The left ST is associated with human speech phoneme sounds (Leaver & Rauschecker, 2010). In addition, amyloid positive and negative populations are associated with the left MT difference (Adamczuk et. al, 2016). The ST are also covered by the AU2 AβN, but the AU2 peaks are located in the transverse temporal gyri that encompasses the primary auditory cortex, highlighting its significance in auditory functions (DeWitt & Rauschecker, 2012). Similarly, as with CC1 the VIS4 AβN covers the IT gyri, but the VIS4 peaks are found in the fusiform cortices, which, in the left hemisphere is related with reading and processing any meaningful visual stimulus (Devlin et al., 2006). The VIS4 AβN also covers the lateral occipital cortices (LOC), which integrates retinotopic and object-selective processing (Sayres & Grill-Spector, 2008). Another AβN, being VIS1 has its most intense point in the LOC, but only in the left hemisphere. VIS1 is also involved with the right lingual cortex (LI) and the right posterior pericalcarine (PEC). Contralaterally, in the left LOC, the VIS2 AβN-peak is found in the right LOC, noting that the VIS2 AβN is dominant to the left hemisphere. The VIS5 AβN peaks in the PEC and covers the LI, being regions covered by VIS1, but VIS5 is bilateral. In addition, VIS5 covers the cuneus cortices (CUN). Similar as VIS1 the VIS3 AβN has its peaks in the LOC, bilaterally. VIS3 also covers CUN, superior parietal cortices (SP), which are critical for sensorimotor integration, by maintaining an internal representation of the body's state (proprioception; Wolpert et al., 1998). The peaks of the SM1 AβN are located in the SP. The SM1 also covers the inferior parietal cortices (IP), which, in the left hemisphere, is a short-term storage of phonological information (Geranmayeh et al., 2012). The peaks of the SM2 AβN are located in the superior precentral cortices (PRC), which controls body movement (Penfield & Boldrey, 1937). In addition, SM2 covers the superior frontal cortices (SF), which are implicated in movement, cognitive control, working memory, among other functions central to decision making and executive control (Zhang et al., 2012; Cañas et al., 2018; Sallet et al., 2013, Boisgueheneuc et al., 2006). The peaks of the SM3 AβN are located in the postcentral cortices (POC), which are related with motor for tongue and larynx (Roux et al., 2018). In addition, SM3 covers the SP and the paracentral cortices (PAC), which are related with blushing (Nikolić et al., 2024). The DM4 AβN is also covered by the SF, but has its peaks in the rostral anterior cingulate, which may resolve emotional conflicts (Etkin et al., 2006). DM4 is also involved with the caudal anterior cingulate cortices. The DM2 AβN peaks in the isthmus of the cingulate cortices and also covers the precuneus cortices (PCUN), which are related with conscious recall of visual memory (Fletcher et al., 1995). The DM3 AβN peaks in the PCUN. The DM1 AβN peaks in the PAC. DM1 also covers the SF. The AU1 AβN peaks in the supramarginal cortices, which, in the left hemisphere, is related with processing phonological inputs and outputs (Oberhuber et al., 2016). The AU1 also covers the POC and the insular cortices, which are related with widely different functions, ranging from pain perception and speech production to the processing of social emotions (Nieuwenhuys et al., 2012). The CC2 AβN peaked in the rostral middle frontal cortices, which are related with schematic control and may be active during sequential tasks (Badre et al., 2019). CC2 also covers SF and the frontal pole cortices, which relate to many aspects of cognition and may process costs and benefits in actions to improve future choices (Tsujimoto et al., 2011). CC3 has its AβN peaks in the SF, but also covers the PRC.

This comprehensive overview elucidates the primary anatomical locations and their functional roles within the DKT framework, offering a clear understanding of how NeuroMark Aβ Networks interact with specific DKT ROIs. Additionally, a note about the DKT ROIs, is that the entorhinal cortex aligns extensively with existing literature and is identified as one of the earliest sites for tau neurofibrillary tangle formation (Lee et al., 2022).
